# Secondary Structure Prediction for RNA Sequences Including N^6^-methyladenosine

**DOI:** 10.1101/2021.04.26.441443

**Authors:** Elzbieta Kierzek, Xiaoju Zhang, Richard M. Watson, Ryszard Kierzek, David H. Mathews

## Abstract

There is increasing interest in the roles played by covalently modified nucleotides in mRNAs and non-coding RNAs. New high-throughput sequencing technologies localize these modifications to exact nucleotide positions. There has been, however, and inability to account for these modifications in secondary structure prediction because of a lack of software tools for handling modifications and a lack of thermodynamic parameters for modifications. Here, we report that we solved these issues for N^6^-methyladenosine (m^6^A), for the first time allowing secondary structure prediction for a nucleotide alphabet of A, C, G, U, and m^6^A. We revised the RNAstructure software package to work with any user-defined alphabet of nucleotides. We also developed a set of nearest neighbor parameters for helices and loops containing m^6^A, using a set of 45 optical melting experiments. Interestingly, N^6^-methylation decreases the folding stability of structures with adenosines in the middle of a helix, has little effect on the folding stability of adenosines at the ends of helices, and stabilizes the folding stability for structures with unpaired adenosines stacked on the end of a helix. The parameters were tested against an additional two melting experiments, including a consensus sequence for methylation and an m^6^A dangling end. The utility of the new software was tested using predictions of the structure of a molecular switch in the MALAT1 lncRNA, for which a conformation change is triggered by methylation. Additionally, human transcriptome-wide calculations for the effect of N^6^-methylation on the probability of an adenosine being buried in a helix compare favorably with PARS structure mapping data. Now users of RNAstructure are able to develop hypothesis for structure-function relationships for RNAs with m^6^A, including conformational switching triggered by methylation.

## Introduction

It has long been appreciated that covalent modification of RNA is used by nature to expand the chemical repertoire of the four common nucleotides. tRNAs, in particular, are known to have prevalent modifications, and the roles of some of these have been elucidated^1^. For mRNAs and long non-coding RNAs (lncRNAs), it had been harder to identify sites of modification until recently when new methods were developed using next generation sequencing technologies to identify modifications^2,3^. Modifications including deamination to inosine^4,5^, pseudo-uridylation^6–8^, 5-methylation of cytosine^9^, and N^6^-adenosine methylation^10–15^ now can now be localized transcriptome-wide.

N^6^-methyladenosine (m^6^A) is considered the most prevalent modification in mRNA, and m^6^A is also widespread in lncRNAs^16,17^. It is known to have writers that apply the modifications to specific positions (methyltransferases including METTL3 and METTL14), readers that identify sequences with N^6^-methylation (RNA-binding proteins including YTHDF2 and the YTH family), and erasers (demethylases including FTO and ALKBH5) that can remove the modification, restoring the base to adenine^18–21^. Furthermore, there are hundreds of sites for which the m^6^A modification consensus site is conserved between the mouse and human genomes^13^. The impacts of N^6^-methylation are being elucidated^22,23^. For example, N^6^-methylation is known to cause structural switches that, for example, can expose protein binding sites that are otherwise not available for binding^24^. Additionally, m^6^A can regulate splicing^25^.

RNA secondary structure prediction is in widespread use to help determine structure-function relationships^26,27^, but has not been generally available for understanding the roles of covalent modifications^28^. For unmodified sequences, secondary structure prediction has been used to identify microRNA binding sites^29^, design siRNAs^30,31^, identify protein binding sites^32^, and discover functional RNA structures^33–35^. These types of calculations have not been able to account for modifications without extensive user intervention because a set of nearest neighbor parameters are needed for estimating the folding stability of structures that include modifications^28,36^. A number of studies have demonstrated an impact on folding stability by modifications^37–42^, but no complete set of parameters have been available for RNA folding, as there are for RNA folding with the four prevalent bases^43^. At the same time, no software has been available for handling a larger alphabet of sequences containing modifications. This led to chicken-and-egg problem; without software, there was no impetus to assemble parameters and without parameters there was no reason to write the software.

In this work, we developed a full set of nearest neighbor parameters for a folding alphabet of m^6^A, A, C, G, and U nucleotides. These parameters account for helix and loop formation, and they are based on optical meting experiments for 32 helices with m^6^A-U base pairs and 13 oligonucleotides with m^6^A in loop motifs. We also modified the RNAstructure software package to accept user-defined folding alphabets and to read and utilize thermodynamic parameters for these extended alphabets^44^. Together, these advances allow the prediction of RNA secondary structures for sequences with m^6^A. We demonstrate, for calculations with human mRNA sequences known to contain m^6^A, that N^6^-methylation alters the folding landscape so that m^6^A is less likely to be buried in a helix, i.e. stacked between two base pairs. We also provide a model for the RNA secondary structures of the methylation-triggered conformational change in the lncRNA metastasis-associated lung adenocarcinoma transcript (MALAT1).

## Results

### Overview of Methods

Secondary structure prediction for RNA sequences including m^6^A requires both a set of nearest neighbor folding parameters and software capable of using the set of parameters. An overview of the methods is illustrated in Figure 1. RNA secondary structure prediction requires both parameters for evaluating folding stability and a search algorithm to identify the optimal structure given the parameters^26,27,45^. In our RNAstructure software, we use nearest neighbor parameters to estimate folding free energy change^36^ and set of dynamic programming algorithms that predict optimal structures^46,47^.

**Figure 1.**
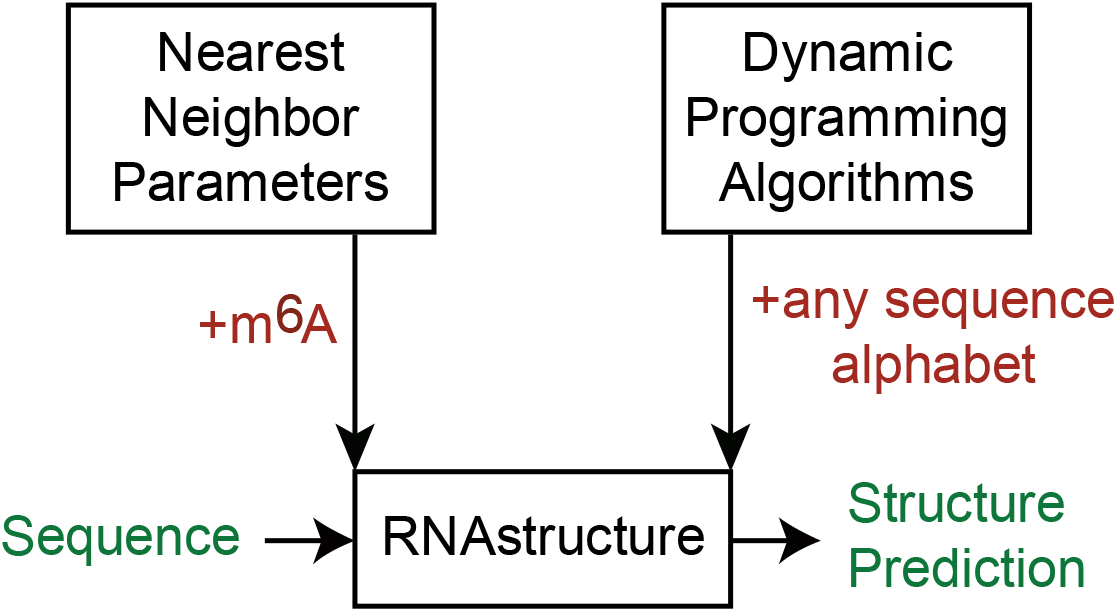
Overview. In this study, we advanced the RNAstructure software package^44^ (at center) to be capable of predicting secondary structures for sequences with the m^6^A nucleotide. RNA secondary structure prediction by RNAstructure relies on nearest neighbor parameters for estimating folding stability and dynamic programming algorithms for estimating structures and base pair probabilities. Here we fitted nearest neighbor parameters for m^6^A to optical melting data and revised the dynamic programming algorithms to be capable of considering any sequence alphabet.

We built a database of optical melting experiments of oligonucleotides including m^6^A and then used linear regression to fit nearest neighbor parameters. We also extended the functionality of RNAstructure^44^ to recognize modified nucleotides in sequences and to use parameters for sequence alphabets beyond the four common nucleotides. The m^6^A modification parameters are the first to take advantage of this new feature.

### Helix Nearest Neighbor Parameters for m^6^A

The full set of Turner nearest neighbor rules for estimating RNA folding stability are based on optical meting experiments of 802 oligonucleotides and use 294 parameters^36,48,49^. We have shown, however, that the precision of a subset of parameters is more important than others for the precise prediction of secondary structure^50^. Following that work, we focused our experiments on estimating parameters for helices, dangling ends, and terminal mismatches.

Our first goal was to fit the 15 stacking nearest neighbor parameters for m^6^A-U pairs adjacent to Watson-Crick pairs, G-U pairs, or m^6^A-U pairs. For this study, 29 fully helical duplexes containing m^6^A-U pairs were synthesized and optically melted. This provides a total database of 32 fully helical duplexes with m^6^A-U base pairs. Table S1 provides the duplexes and the stabilities determined by optical melting. These specific oligonucleotide sequences were chosen, in part, because analogous model RNA helices with A in the m^6^A position had been previously studied by optical melting (with the exception of GGUUAACC_2_). This allows us to directly compare the folding stability with and without N^6^-methylation. We calculated the change in folding stability (ΔΔG°_37_) per methylation as compared to the unmethylated duplex. Figure S1 shows that the ΔΔG°_37_ is highly dependent on the adjacent sequence, ranging from +2.1 to −0.1 per methylation where positive free energies are destabilizing for methylation. Therefore, to estimate folding stabilities for duplexes with m^6^A-U pairs, a full nearest neighbor model is needed to account for the sequence dependency.

Linear regression was used to fit the nearest neighbor parameters for folding free energy change. Figure 2A shows the increments in comparison to the same stack with A-U pairs and Table S2 provides the values. The free energy changes range from −1.79±0.25 kcal/mol to +1.45±0.57 kcal/mol. As expected based on prior optical melting experiments for duplexes with m^6^A-U pairs^37,39^, nearest neighbor stacks for methylated A-U pairs are less stable than stacks for unmethylated A-U pairs. On average, the stacks with m^6^A-U pairs are 0.4 kcal/mol less stable per methylation. There are exceptions, however; an m^6^A-U pair followed by a U-A pair is as stable as an A-U pair followed by a U-A pair (−1.10 kcal/mol). The most unstable stack has two m^6^A-U pairs. Like A-U pairs, when the m^6^-U pair is adjacent a G-C it is more stable than when adjacent to A-U. Also like A-U pairs, m^6^A-U pairs adjacent to G-U are less stable than those adjacent to A-U pairs.

**Figure 2.**
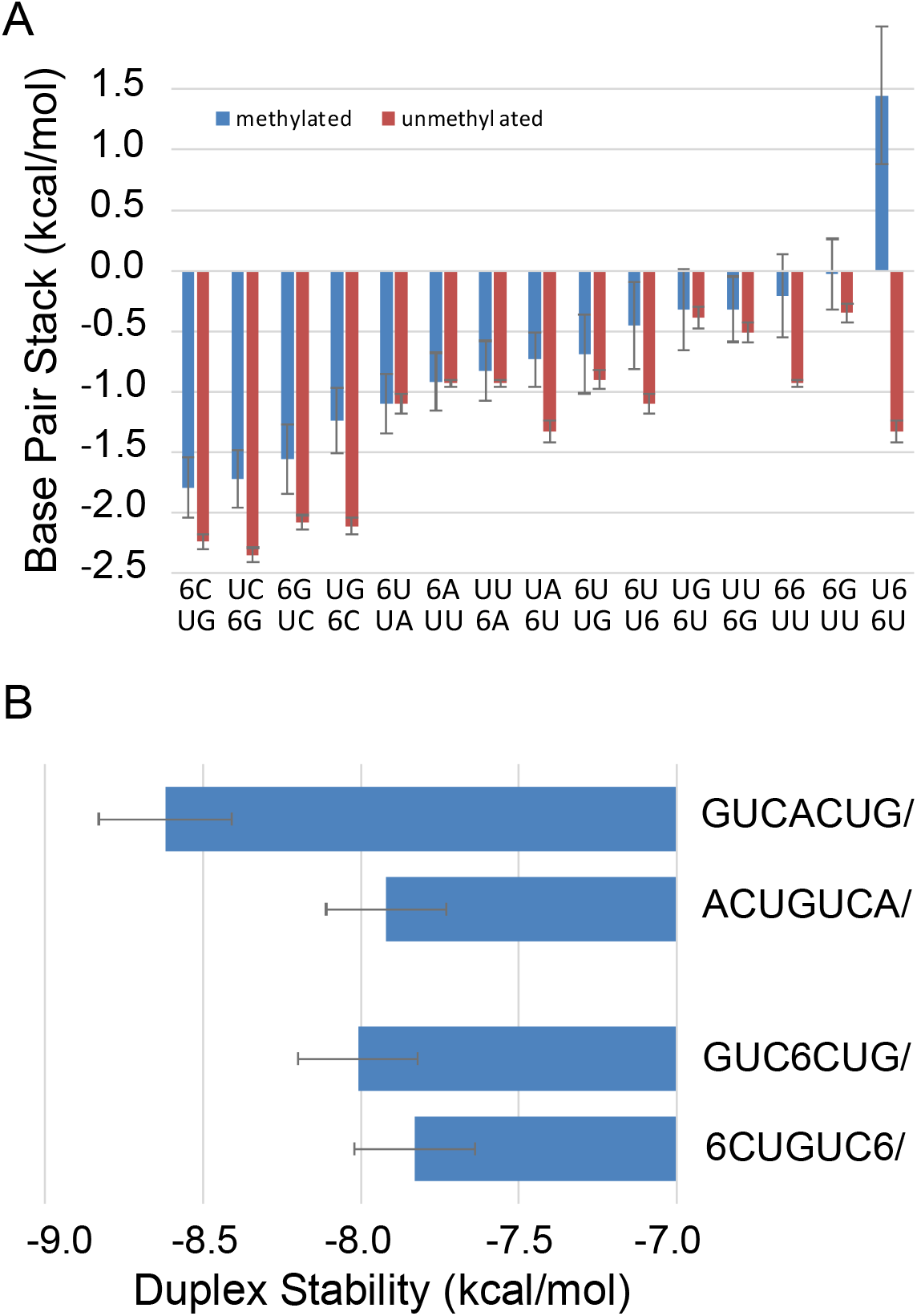
**Panel A. The nearest neighbor parameters for helix stacks**. The position of the m^6^A is indicated by 6. The stacking parameters are compared for methylated (blue; i.e. m^6^A-U base pairs) and unmethylated (red; i.e. A-U base pairs) sequences for analogous nearest neighbors. The unmethylated stacks (i.e. A-U base pairs) are those of Xia et al.^51^ for adjacent Watson-Crick pairs and those of Chen et al.^73^ for adjacent G-U pairs. Stacks with m^6^A-U pairs are generally less stabilizing than analogous stacks with A-U pairs. **Panel B. Terminal m**^**6**^**A-U pairs are not destabilizing**. The top two sequences (Watson-Crick paired with a complementary strand) have the same nearest neighbor stacks, but the second helix has two terminal A-U pairs^51^. This costs 0.7 kcal/mol of stability. The bottom two sequences also have the same nearest neighbor stacks, but the second has two terminal m^6^A-U pairs. Here the stability cost is 0.18 kcal/mol and not outside of the uncertainty estimate. On average, terminal A-U pairs cost 0.45 kcal/mol of stability^51^, but terminal m^6^A-U pairs are not destabilizing.

An unexpected feature of terminal m^6^A-U pairs is that they require no terminal penalty, although terminal A-U pairs receive a +0.45 ± 0.04 kcal/mol penalty per A-U pair at the end of a helix^51^. Two findings support this. First, when a terminal parameter is included as a parameter in the linear regression fit, the value is +0.13 ± 0.17 kcal/mol, which is not significantly different from 0 kcal/mol. Second, our dataset includes two helices with the same nearest neighbor stacks, but with different helix ends (Figure 2B). Previously, it was noted that this pair of helices, when unmethylated, had markedly different stability (0.70±0.28 kcal/mol), with the helix with A-U ends less stable^51^. For the methylated helices, the difference is small (0.18±0.27 kcal/mol). This demonstrates that a terminal m^6^A-U base pair has overall similar stability to a terminal A-U base pair because a terminal A-U pair has a more favorable stack but requires the terminal A-U penalty.

### Loop Nearest Neighbor Parameters for m^6^A

For secondary structure prediction, parameters need to also be extrapolated for loop formation. The stability of a 3’ dangling m^6^A had been previously measured^37^. Additional optical melting experiments were performed for two m^6^A 3’ dangling ends, an m^6^A 5’ dangling end, and seven terminal mismatches involving at least one m^6^A. One hairpin loop was measured with an m^6^A in the loop and not adjacent to the helix end. One 2×2 internal loop was measured with symmetric tandem G-m^6^A pairs. The loop sequences were chosen such that analogous sequences with A instead of m^6^A had been previously studied, so that the effect of methylation on stability can be quantified. Table S3 provides the measured stabilities for these model structures and Table S4 shows the stability of the loop motif in comparison to the motif with A.

As shown by Figure 3, an m^6^A as a dangling end or as a component in a terminal mismatch stabilizes secondary structure formation to a greater extent than an analogous A. On average, the m^6^A dangling end is −0.43±0.15 kcal/mol more stable than the analogous A dangling end for the 3’ and 5’ dangling ends studied here. Terminal mismatches for m^6^A-m^6^A, G-m^6^A, m^6^A-G, and m^6^A-C on Watson-Crick or G-U terminal pairs are on average −0.28±0.26 kcal/mol more stabilizing than the analogous A-A, G-A, A-G, or A-C terminal mismatches. This stabilizing effect is sequence dependent; the ΔΔG°_37_ ranges from −0.74 kcal/mol (G-m^6^A mismatch on a U-G pair) to +0.02 kcal/mol (G-m^6^A mismatch on an A-U pair). An m^6^A-m^6^A mismatch on an m^6^A-U pair is more stable than the m^6^A-m^6^A mismatch on an m^6^A-U pair by −0.42±0.40 kcal/mol.

**Figure 3.**
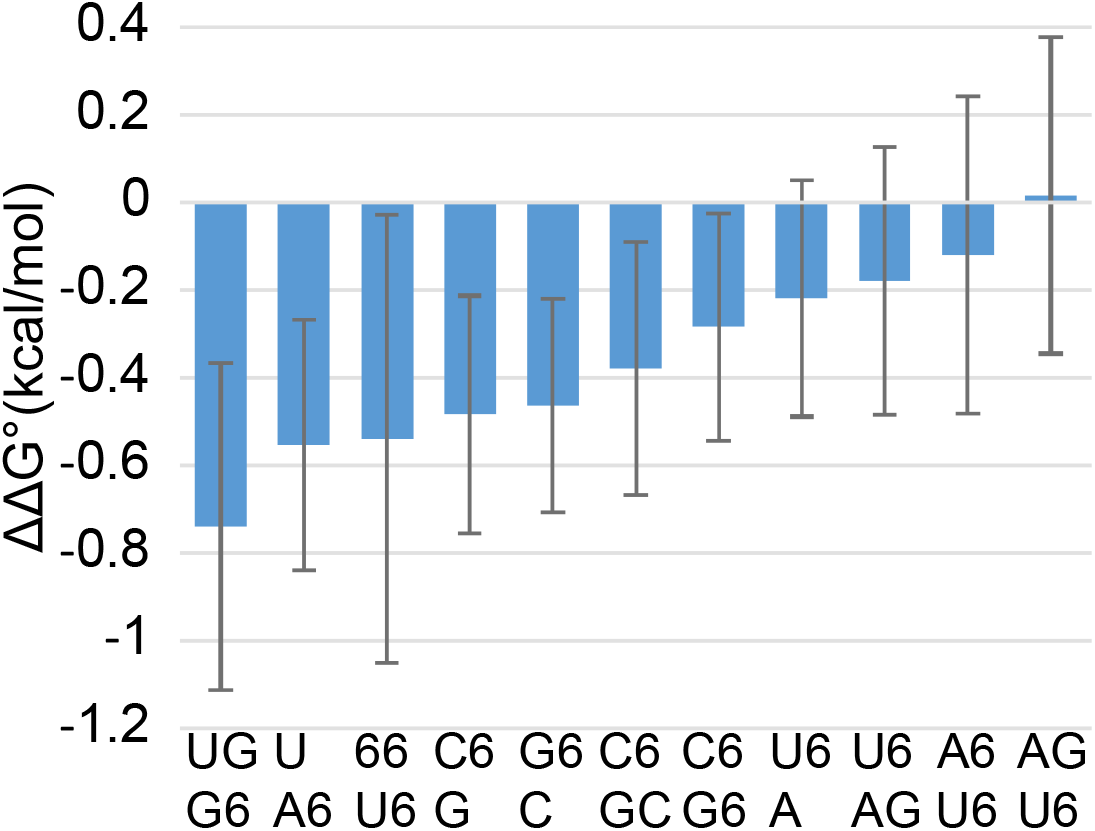
m^6^A Stacking on a Helix End Stabilizes Secondary Structure as Compared to A Stacking. The ΔΔG°_37_ (kcal/mol) for dangling ends and terminal mismatches as a result of N^6^-methylation (Table S4) is shown, where negative values mean greater folding stability for m^6^A than A. The motifs shown here have a terminal base pair (left side of motif), and either a dangling end or terminal mismatch right (right side of motif). On average, the methylated motifs are more stabilizing than the unmethylated motifs, although the extent of the stabilization is sequence dependent.

The hairpin loop structure with m^6^A is marginally less stable than the analogous hairpin loop with A (ΔΔG°_37_ = 0.23±0.24 kcal/mol; Table S4). The 2×2 internal loop with tandem G-m^6^A pairs is also marginally less stable than the analogous loop with tandem G-A pairs (ΔΔG°_37_ = 0.33±0.53 kcal/mol; Table S4). Both stability changes are within the uncertainty estimates, suggesting that they are not substantial differences.

### Additional Experiments to Test the Parameters

To test our parameters, we performed additional melts of duplexes. The first is a duplex with all base pairs, incorporating a consensus N^6^-methylation site, GGACU, where we determined the helix stability with and without methylation. The second is an additional 3’ dangling m^6^A to test our assumption that dangling m^6^A are stabilized by −0.3 kcal/mol compared to dangling A. Table S5 provides the stabilities determined by optical melting and Table S6 shows how well the stabilities are estimated with our nearest neighbor parameters.

We conclude from these tests that the nearest neighbor parameters are accurate enough to be used for RNA secondary structure prediction^48,50^. The estimates for the duplex stabilities are within the uncertainties propagated for the experiment and the nearest neighbor parameters (ΔΔG°_37_ column of Table S6). The unmethylated consensus duplex is estimated by nearest neighbor parameters to be more stable (by −0.48±0.73 kcal/mol) than it is by experiment. The methylated consensus duplex is estimated by nearest neighbors to be less stable than it is (by 0.85±0.97 kcal/mol). These deviations are 2.9% and 5.5% of the experimentally determined values. The estimated stability of the duplex with the dangling m^6^A closely matches the experimental value (ΔΔG°_37_ of 0.01±0.84 kcal/mol).

### RNAstructure Software Modifications

To predict RNA secondary structures for sequences with A, C, G, U, and m^6^A, we modified the command line programs in the RNAstructure software package to accept extended alphabets of nucleotides^44^. By default, the software interprets sequences as standard RNA, but a command line switch can specify an alternative alphabet. For example, the nearest neighbor parameters for a DNA alphabet composed of A, C, G, and T has long been available. Now, because of this work, the nearest neighbor parameters for an RNA m^6^A alphabet is available.

The key to an extended alphabet is the specification of the nucleotides and pairs (Figure S2). A common architecture across the RNAstructure programs means that the command line programs are capable of using the extended alphabets, which can include any number of characters. This includes the prediction of minimum free energy structures, base pair probabilities, maximum expected accuracy structures, and folding stability for structures. Each nucleotide must be encoded by a single-character, and we chose “6” or “M” as the character to encode m^6^A in sequences and in the m^6^A nearest neighbor parameter tables. The Methods section details our estimates for the m^6^A nearest neighbor parameters.

Nearest neighbor parameter tables are read from disk as programs start. Each parameter table requires additional rows and columns to provide the nearest neighbor parameters values for those nucleotides, although the dimensionality of the tables stays the same. For example, a base pair stack table is four-dimensional because the sequence of four positions is required to estimate the stacking stability of two pairs. When m^6^A is included with RNA, the size of each dimension is increased to five from four. The largest table is the 2×2 internal loop lookup table^36^, which is eight dimensional because it includes the sequence of the two closing base pairs.

### Modeling Conformational Changes as a Result of Methylation

It has been established that N^6^-methylation can alter RNA structure, and because of this m^6^A is considered a conformational switch. To test our new m^6^A nearest neighbor parameters and software, we made a quantitative prediction for the switching of the structure in the lncRNA MALAT1 that opens a binding site for heterogeneous nuclear ribonucleoprotein C (HNRNPC). This has been characterized by Tao Pan and co-workers in an *in vitro* system with a single stem-loop structure^52^. Filter binding experiments demonstrated that the methylated RNA is more accessible to protein binding than the unmethylated RNA. Additionally, enzymatic cleavage by RNase S1, which has specificity for loop regions of RNA, demonstrated increased cleavage 5’ and 3’ to the methylated A, supporting a conformational change.

We used RNAstructure to predict the secondary structure of the 32 nucleotide RNA using stochastic sampling of the structures from the ensemble^53^. This can be used to characterize the structures of RNAs that can fold to more than one structure at equilibrium. We found two predominant structures in the ensemble, as demonstrated in Figure 4A. One of the two structures is that of the stem-loop that was previously predicted and has three of the five nucleotides at the HNRNPC site base paired^52^. Interestingly, the second major structure in the ensemble has two stem-loops and the HNRNPC site is more exposed for protein binding (a single U at the 5’ end of the binding site is base paired). RNAstructure estimates a shift in the population from the closed (protein-occluded) structure to the open (protein-accessible) structure in agreement with the experimentally measured shift in protein binding. In the absence of methylation, the ratio of closed:open is estimated to be 62:38, but in the presence of N^6^-methylation the ratio is estimated to be 41:59. This demonstrates a quantitative prediction of shift in ensemble folding behavior with methylation that explains how the methylation accomplishes the structural switching.

**Figure 4.**
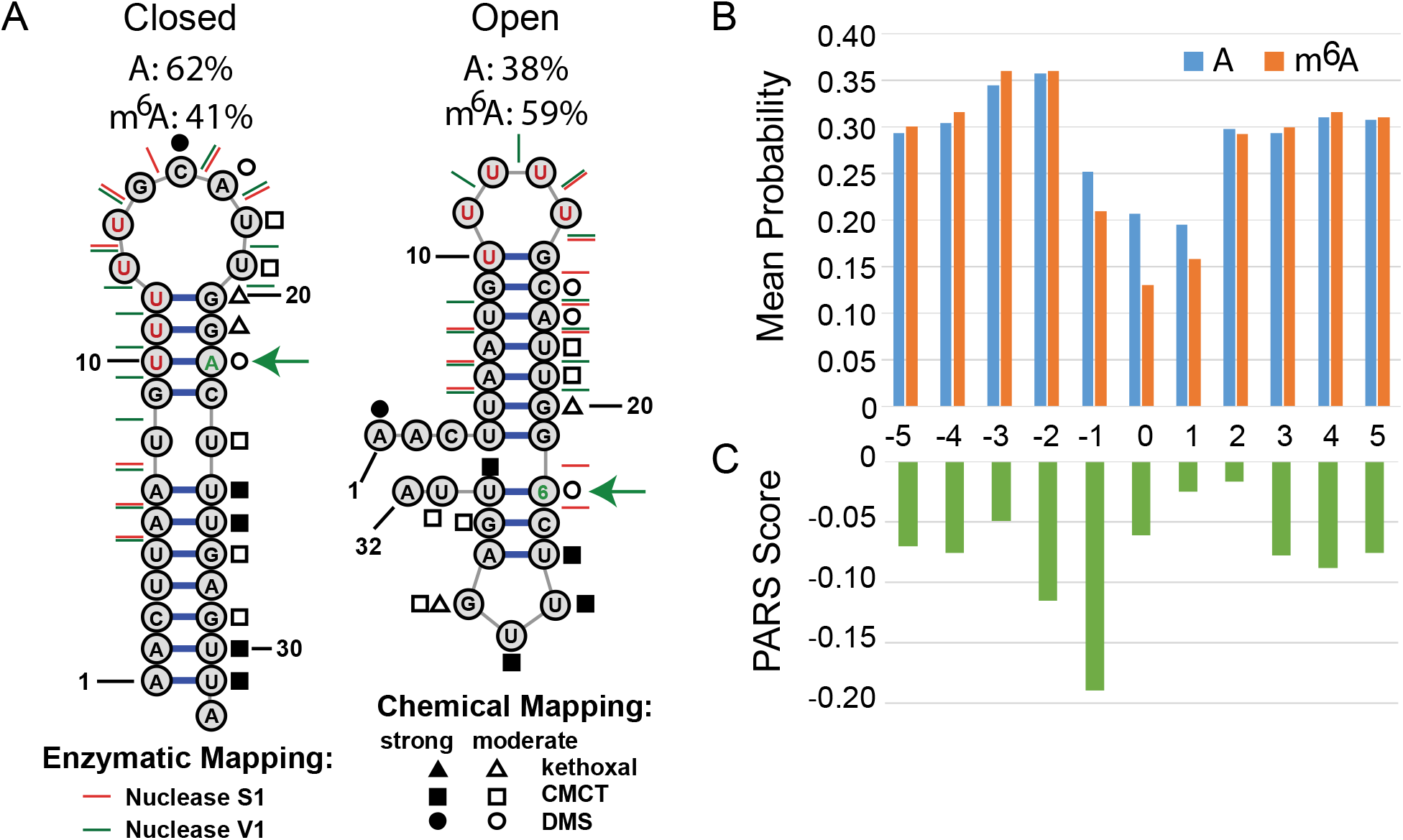
Tests of the new m^6^A nearest neighbor parameters and RNAstructure software. **Panel A. The Conformational Switch of MALAT1 RNA**. Secondary structure was predicted for the model methylation-activated switch for HNRNPC binding. In the absence of methylation, the left (closed) structure is predicted to predominate the ensemble, where the HNRNPC protein binding site (marked in red nucleotides) is occluded to binding. When A22 is methylated to m^6^A, the right (open) structure is predicted to predominate, where the HNRNPC binding site is more accessible. In the absence of methylation, the ratio of closed:open is estimated to be 62:38. This estimate switches to 41:59 for the methylated sequence. Chemical and enzymatic probing results are superimposed on the structures. Mapping data for the unmethylated sequence are superimposed on the closed conformation (left) and mapping data for the methylated sequence are superimposed on the open structure (right). The chemical agents act on Watson-Crick faces and prefer loop nucleotides, although they also act on helix ends and G-U pairs^54^. Nuclease S1 prefers loop regions and Nuclease V1 prefers helical regions^54^. The chemical and enzymatic mapping data support a mixture of the two structures both with and without N^6^-methylation. **Panel B. The Average Probability that A or m**^**6**^**A are Buried in a Helix at the Position of a High-Confidence m**^**6**^**A sites in the Human Transcriptome**. The mean probability that an A or m^6^A is base paired and stacked between two adjacent pairs for 18,026 sites of N^6^-methylation, as estimated by RNAstructure. Position 0 is the site of methylation. N^6^-methylation is estimated to further open the structure at the methylation site. **Panel C. The Average PARS Scores for Accessibility for the 18**,**026 sites of N**^**6**^**-methylation in the Human Transcriptome**. Lower PARS scores indicate higher counts of nuclease S1 cleavage relative to nuclease V1 cleavage and therefore a higher likelihood of being unpaired. The RNAstructure predictions and the PARS data both show considerable single-stranded character at the site of N^6^-methylation.

Furthermore, we probed the 32 nucleotide RNA (extended with a 3’ structural cassette) by chemical mapping with CMCT (1-cyclohexyl-3-(2-morpholinoethyl) carbodiimide metho-p-toluenesulfonate), DMS (dimethyl sulfate), and kethoxal (Figure 4A). The chemical mapping data and the prior enzymatic mapping^52^ data are consistent with the sequence populating more than one structure; neither of the two proposed structures alone fully explains the data. Nuclease S1 prefers to cut in loop regions and Nuclease V1 prefers to cut in helical regions^54^, but a number of cleavages occur in tandem at the same phosphodiester bond. This indicates a mixture of structures. The loss of V1 cleavages between U10 and U11 supports an increase in the population of the open structure upon methylation. The S1 cleavage that emerges 5’ and 3’ to m^6^A22 upon methylation is consistent with the increase in population of the open structure. The chemical mapping data also suggest a mixture of two structures; the three agents used react with moieties on the Watson-Crick faces and paired bases are generally more protected than unpaired bases^54^. A number of base reactivities support a population of the closed structure for both sequences, including C16, A17, U19, and U20. The CMCT reactivities at U26 and U30 support the second hairpin loop in the open structure, both with and without methylation. The loss of reactivity upon methylation at C15 and the gain of kethoxal reactivity upon methylation at G29 are consistent with the shift towards the open conformation upon methylation. In summary, the structure predictions with m^6^A parameters provide a more complete picture of conformational switching, which is more complicated than a simple on-off switch and better reflected as a shift in the Boltzmann-weighted ensemble between conformations.

### Transcriptome-Wide Predictions with m^6^A

To further test our new m^6^A nearest neighbor parameters and software, we predicted structures for 18,026 mRNAs that were identified as having N^6^A methylation by whole transcriptome sequencing^55^ and for which PARS structure mapping data are available^56^. We used the nearest neighbor parameters and RNAstructure package to estimate the probability that the methylation site is buried in a helix, i.e. in a base pair stacked between two other base pairs, for both the unmethylated and methylated sequence (Figure 4B). We used 800 nucleotide fragments of local sequence to estimate the pairing probability because we previously found that pairing probability estimates for 800 nucleotide fragments reasonably match those for global secondary structure prediction^30^. This is a reasonable balance between accuracy and total calculation time.

We find that the unmethylated A at the methylation site is less likely to be buried in a helix than adjacent nucleotides (Figure 4B). This is intuitive because adjacent nucleotides at the consensus site are often G or C, and A is more predominant in loops in RNAs with known structure^57^. There is a substantial shift in the probability of m^6^A being buried in a helix relative to A (21% for A and 13% for m^6^A). This suggests there could be widespread structural switching being affected by N^6^-methylation. We can also compare our results to PARS data for the same sequences (Figure 4C)^39,56^. A PARS score quantifies the enzymatic cleavage estimate of local pairing and the experiment is performed transcriptome wide. A lower PARS score indicates greater nuclease S1 cleavage relative to nuclease V1 and thus a greater extent of unpairing because nuclease S1 has specificity for loops and nuclease V1 has specificity for helices^54,58–60^. The PARS scores at the methylation site also demonstrate a propensity to be unpaired at the methylation site, but the minimum average PARS score is at the nucleotide 5’ to the m^6^A site. A possible explanation for the discrepancy is that PARS attributes S1 cleavages to the base 5’ to the cleavage site, assuming that the base 5’ to the cleavage is unpaired. Cleavage can also occur when the base 3’ to the cleavage site is unpaired and therefore the PARS scores 5’ to the methylation site might be overestimating the propensity of being unpaired, in that some of the propensity of being unpaired should be attributed to the methylation site. For example, the prior S1 mapping of 5S rRNA structure is consistent with cleavages both 5’ and 3’ to unpaired nucleotides (Figure S3)^61^. Prior analysis of PARS scores for methylation sites also concluded that the data indicate the m^6^A is positioned in structures at the transition between base paired regions and loop regions, consistent with our structure prediction estimates^39^.

## Discussion

Here we provide the first complete nearest neighbor model for a folding alphabet including modified nucleotides. Because m^6^A is considered the most abundant modification in mRNA and is known to affect folding stability, we chose m^6^A as the first modification to study. The full nearest neighbor model for secondary structure prediction requires both helical stack parameters and also loop parameters. We know from a sensitivity analysis of secondary structure prediction that, for loops, accurate parameters are most important for dangling ends and terminal mismatches^48,50^, accordingly we focused our experimental effort on these motifs. We also observed marginal differences in stability for hairpin and internal loops containing m^6^A as compared to the same sequences without the N^6^-methylation. Subsequent studies could be focused on understanding and modeling folding stability differences for loops with m^6^A.

The other component of this study was advancing RNAstructure to work with sequences with nucleotides beyond A, C, G, and U. We provide command line tools that are ready to make quantitative predictions of structure and folding stability for sequences with m^6^A. Given the software, we plan expand our work in the future to include alphabets with inosine and pseudouridine. Both have helical nearest neighbor parameters available for stacks on Watson-Crick pairs^40,41,62^, and both could be extended to full nearest neighbor parameters sets with additional optical melting experiments.

The two loops studied here with N^6^-methylations both had marginally less folding stability than the analogous unmethylated loops. Solution structures are available for each of the A-containing loops, and these structures provide clues as to why the stabilities would be only marginally changed by methylation. The hairpin loop, GGC<u>GUAAUA</u>GCC, has the first A in the loop (A6; the site of our m^6^-methylation) stacked at the apex of the loop on the adjacent A (A7)^63^. Because A6 is not hydrogen bonding in the structure, a methylation at N^6^ can be accommodated in the preferred syn orientation by the structure without change^64^. For the internal loop with tandem G-A pairs, the pairs are trans-sugar-Hoogsteen pairs, i.e. the N^6^ position of the A is hydrogen bonded with the G at the N^3^ position^65^. For each methylated A, one hydrogen of N^6^ is available to form this hydrogen bond, placing the methyl in the preferred syn orientation^64^. However, the second hydrogen of A N^6^ is close to O4’ of the G (ranging from 2.34 to 3.36 Å in the 15 deposited NMR models). This suggests the structures would need at least small changes to accommodate the syn methyl to avoid a steric clash^39,64^.

Recent studies demonstrated the ability of computational methods to estimate folding free energy changes^66–72^. In this work, we performed optical melting experiments to determine the folding stabilities of small model systems with m^6^A and fit nearest neighbor parameters to these data. Future work, however, could rely on computation or a mixture of computation and experimentation. Hopfinger et al., for example, estimated helical stacking nearest neighbor parameters for the eight stacks with m^6^A-U pairs adjacent to Watson-Crick pairs^66^. Overall the agreement of their estimates against our experimental values is excellent, with a root mean squared deviation of 0.30 kcal/mol. The largest single deviation is for a U-m^6^A pair followed by a G-C pair, where their estimate overstabilized the stack by 0.6 kcal/mol (Figure S4). Loop folding stabilities continue to be more of a challenge to estimate using computational methods because the conformational flexibility requires extensive sampling^71^.

With this work, we demonstrate the position of m^6^A in a structure determines whether folding stability is increased, decreased, or unchanged relative to the same structure with A. It was previously known that N^6^-methylation of an A-U pair in the middle of a helix would decrease the helix folding stability^37,39^. Our stacking parameters now quantify this sequence-dependent change (Figure 2A). It was also previously known that m^6^A stacking on the end of a helix would stabilize the helix more than an analogous A. In this work, we also discovered that an m^6^A-U base pair at the terminal position of a helix provides roughly the same folding stability as an analogous A-U base pair. This is because terminal A-U pairs destabilize helices with a penalty of +0.45 kcal/mol^51^ that is not needed for terminal m^6^A-U base pairs (Figure 2B). Recently, it was also discovered that terminal G-U base pairs in helices do not need an end penalty^73^. These results, taken together, show why N^6^-methylation is a potent switch of secondary structure.

Our transcriptome-wide calculations also suggest that structure switches from N^6^-methylation might be widespread (Figure 4B). It will be interesting to perform similar calculations with other widespread covalent modifications, such as inosine. There is potential to identify structural mechanisms by which covalent modifications exert changes in protein binding, transcript stability, or gene expression.

## Acknowledgements

This work was supported by National Institutes of Health grant R01GM076485 to D.H.M. and by National Science Center grants: UMO-2020/01/0/NZ6/00137 to E.K. and UMO-2019/33/B/ST4/01422 to R.K.

## Methods

### Synthesis of Oligonucleotides with m^6^A

Oligoribonucleotides were synthesized on a BioAutomation MerMade12 DNA/RNA synthesizer using β-cyanoethyl phosphoramidite chemistry and commercially available RNA phosphoramidites (ChemGenes, GenePharma) and protected N^6^-methyladenosine phosphoramidite, which was synthetized according to a standard protocol. (Synthesis of N^6^-methyl adenosine via Dimroth rearrangement followed by protection of the 5’-hydroxyl with dimethoxytrityl and 2’-hydroxyl with tert-butyldimethylsilyl. Next, 5’- and 2’-protected N^6^-methyladenosine was treated with 2-cyanoethyl N,N,N′,N′-tetraisopropylphosphorodiamidite^74,75^.) Oligoribonucleotides were deprotected with aqueous ammonia/ethanol (3/1 v/v) for 16 h at 55°C. Silyl protecting groups were cleaved by treatment triethylamine trihydrofluoride. Deprotected oligonucleotides were purified by silica gel thin layer chromatography (TLC) in 1-propanol/aqueous ammonia/water (55/35/10 v/v/v) as described previously^51,75^.

### Optical Melting Data

The thermodynamic measurements were performed for nine various concentrations of RNA duplex in the range of 0.1 mM – 1 µM in buffer containing 1 M sodium chloride, 20 mM sodium cacodylate, and 0.5 mM Na_2_EDTA, pH 7. Oligonucleotide single strand concentrations were calculated from the absorbance above 80°C and single strand extinction coefficients were approximated by a nearest-neighbor model^76^. Absorbance vs. temperature melting curves were measured at 260 nm with a heating rate of 1°C/min from 0 to 90°C on JASCO V-650 spectrophotometer with a thermoprogrammer. The melting curves were analyzed and the thermodynamic parameters calculated from a two-state model with the program MeltWin 3.5^77^. For most model RNAs, the ΔH° derived from T_M_^-1^ vs. ln(C_T_/4) plots is within 15% of that derived from averaging the fits to individual melting curves, as expected if the two-state model is reasonable.

### Linear Regression

Linear least-squares fitting to determine RNA stacking stabilities was performed with a custom Python program using the statsmodels ordinary least-squares class (OLS)^78^. For each duplex, to determine the stabilities to be fit, the fixed terms were subtracted, including the stability of base pair stacks with Watson-Crick and G-U pairs only, the duplex initiation term, the terminal A-U penalty term (when needed), and the symmetry term (when needed). The fit was excellent, with coefficient of determination, R^2^, of 0.984. Uncertainty estimates (Figure 2A and Table S2) are the standard errors of the regression. Table S7 shows the stability to be fit and the estimate of the fit. Table S8 shows the number of occurrences of each stacking parameter in the set of fit helices.

### Loop Motif Stability Calculations

Loop motif stabilities (Table S4) are calculated by subtracting the helical component of stability.

For the dangling ends and terminal mismatches, twice the stability increment of the motif is determined by subtracting a reference helix stability from the stability of the duplex with the motif^79^:

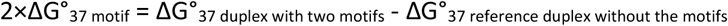

The factor of two is present because the self-complementary duplexes have two instances of the motif.

For the hairpin loop, the stability of the loop motif is determined by subtracting the stability of the helical stacks (estimated with nearest neighbor parameters) from the total stability^80^:

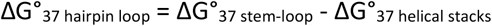

The total helical stack stability is reported as the Reference ΔG°_37_ in Table S4.

For the internal loop, the stability is the total stability of the duplex minus the helical stacks (estimated with nearest neighbor parameters) and minus the stability cost of symmetry (because the duplex is self-complementary)^81^:

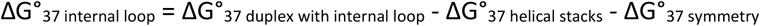

The Reference ΔG°_37_ reported in Table S4 is the sum of the helical stacks and symmetry free energy increments.

### Error Propagation

To estimate uncertainties in free energies (σ), we propagate uncertainty estimates for experiments and nearest neighbor parameters using the standard method for uncorrelated parameters:

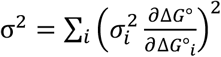

where ΔG°_i_ is the i^th^ term and σ_i_ is the uncertainty in the i^th^ term^50,82^. For the sum of terms used here, this simplifies to:

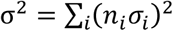

where n_i_ is the number of occurrences of the i^th^ parameter.

For uncertainty estimates for optical melting experiments, we use 4% of the magnitude of the ΔG°_37_. This was chosen as a conservative estimate of the precision of optical melting by Xia et al.^51^. It is twice the mean difference in free energies determined using the two fit methods for optical melting data (Average of Curve Fits and Analysis of T_M_ Dependence) for a database of optical melting experiments.

### Nearest Neighbor Parameter Determination

Nearest neighbor parameters were developed to estimate the folding stability (ΔG°_37_) of sequences with A, C, G, U, and m^6^A. Nearest neighbor parameters are inherited from the 2004 Turner Rules^36^, where a summary of their derivation can be found in Zuber et al.^48^ and examples for their use are available on the Nearest Neighbor Database (NNDB)^43^ website. Helical stacking tables are from Xia et al.^51^ for Watson-Crick stacks and from Chen et al.^73^ for stacks that contain G-U pairs, supplemented with the new stacks determined for m^6^A-U pairs in this work. Following Chen et al., terminal G-U base pairs in a helix are not penalized.

Dangling end m^6^As are stabilized as compared to the analogous A dangling end by the mean additional stability found here (−0.4 kcal/mol). Dangling ends on m^6^A-U pairs are assumed to be the same stability as dangling ends on A-U base pairs. When the stability is measured by an experiment, the measured value is used (Table S4).

Terminal mismatches involving m^6^A are estimated to be more stable than the analogous A terminal mismatch by the mean value found for the terminal mismatches in this study (−0.3 kcal/mol). m^6^A-m^6^A terminal mismatches receive only −0.3 kcal/mol additional stability. A terminal mismatch on an m^6^A-U pair is also stabilized by −0.3 kcal/mol compared to the analogous mismatch on an A-U pair. These effects are additive; an m^6^A-containing terminal mismatch on an m^6^A-U pair receives an additional −0.6 kcal/mol stability than the analogous terminal mismatch with all A parameters. When the stability is measured by an experiment, the measured value is used (Table S4).

Hairpin, internal, and bulge loop initiation costs are length-dependent^36^. The same length-dependent costs are used here, which assumes that m^6^A does not alter the initiation costs.

1×1, 2×1, and 2×2 internal loop stabilities are stored in lookup tables. The stabilities for loops with unpaired m^6^A are taken from the analogous loops with A. And m^6^A-U-closed loops are taken from analogous A-U-closed loops with one change. A-U-closed loops have a 0.7 kcal/mol stability penalty per closure^36^; for m^6^A-U-closed loops, this cost has been removed compared to the analogous A-U-closed loop. Larger internal loops use a terminal stacking table to assign a stability increment for the sequence of the closing pair and first mismatch. Separate tables are used for loops of size 1×n, 2×3, and (>2)×(>2)^36^. These terminal stack tables use the analogous A parameter for stacks with m^6^A. The one exception is that the +0.7 kcal/mol internal loop A-U pair closure penalty is removed for m^6^A-U closures.

Hairpin loop tables for triloop, tetraloops, and hexaloops are unchanged. These tables include stabilities for specific hairpin sequences known by experiment to not be well predicted using nearest neighbor rules^36^. Other hairpin loops are estimated with the sum of a terminal mismatch and a length-dependent initiation. The terminal mismatches for m^6^A use the analogous A parameter.

Multibranch loop initiation parameters are from an experimental fit using a simple linear model^83,84^. The ersatz functional form was found to perform well in a study testing alternative functional forms^85^. Coaxial stacking is included in multibranch and exterior loops^36^. Coaxial stacking between two adjacent helices is assumed to be as stable as a helical stack. For coaxial stacks with an intervening mismatch, there are two stacks. The coaxial stacking increment for the stack where the backbone is not continuous was previously found to be independent of sequence^86^, and the sequence-independent value is used here for stacks involving one or more m^6^As. The other stack is identical to the terminal mismatch stack table.

### Extended Alphabet Implementation in RNAstructure

RNAstructure is a software package written in C++, with a C++ class library that is also wrapped using SWIG to be available to JAVA or Python programs^44^. It is open source and provided for free under the GNU GPL license version 2 at https://rna.urmc.rochester.edu. A number of the command line programs have been updated to be capable of using extended alphabets, including Fold^87^ (secondary structure prediction by free energy minimization), efn2^88^ (estimation of folding free energy changes for secondary structures), partition^47,89^ (partition function calculations for estimating pair, motif, or structure probabilities). A number of programs that rely of the partition function calculations are therefore also able to consider extended alphabets, including AllSub^90^ (prediction of all low free energy structures within an increment of the lowest), design^91^ (design of a sequence to fold to a specific secondary structure), EnsembleEnergy (calculation of the ensemble folding free energy change), MaxExpect^84,92^ (prediction of maximum expected accuracy structures), ProbKnot^93^ (prediction of structures that can include pseudoknots), ProbScan^94^ (estimation of motif probabilities), and stochastic^53^ (stochastic sampling from the Boltzmann ensemble).

The command line tools read the thermodynamic parameters at startup. The switch --alphabet is used to specify the set of parameters to be used. The default is “rna”, the current (2004) Turner rules for estimating RNA folding free energy changes^36,51^. Included with the latest RNAstructure release (version 6.3) is also “m6A”, the parameters discussed here, and “dna”, a set of nearest neighbor rules for DNA secondary structure prediction. The files are a plain text format that was updated (in version 6.0) for extended alphabets. The specification file (Figure S2) is read first, and defines the alphabet and base pairs. Dynamic memory allocation is used to provide the memory needed to store the tables. The parameters themselves are then read from the files.

The 2004 Turner rules gave a terminal base pair penalty for any base pair (A-U or G-U) at the end of a helix that contained a U^36,51,88^. In this work, we found that terminal m^6^A-U pairs did not require this terminal base pair penalty. Additionally, the revised G-U parameters^73^, used with the m^6^A parameters we derived, do not require a terminal base pair penalty. Therefore, we changed the implementation of the energy function to account for this change.

### MALAT1 Calculations

The secondary structure of the 32-nucleotide fragment of MALAT1 was predicted with and without N^6^-methylation of A22 using the stochastic program in RNAstructure^44^. The ensemble size was set to 100 structures. The predicted ensembles show fluctuations in pairs around the two predominant structures, as expected. As examples, terminal base pairs for helices are variably present and the lower helix in the Open structure of Figure 4A can be absent in the Open structure. To classify each sampled structure as Open or Closed, the hamming distance on base pairs was calculated to each the Open and Closed conformation, and the structure was assigned to the conformation with lower distance.

### Chemical mapping of RNA and data analysis

DMS (to modify adenosine and cytidine), CMCT (to modify uridine and guanosine) and kethoxal (to modify guanosine) were used to chemically map secondary structure of 32 nucleotide RNA (with a 3’-structural cassette). The RNA (5’AACUUAAUGUUUUUG*C*AUUGGACUUUGAGUUA<u>CCUUCCGGGCUUCGGUCCGGAAC</u>) was synthesized using the phosphoramidite method on a MerMade synthesizer, deprotected and purified on a 12% denaturing gel. The RNA contained a structure cassette at the 3’ end (underlined), which was designed using RNAstructure to fold independently and allow readout of whole structure of studied RNA^95^. The RNA contained C16-2’-OMe instead of a standard C nucleotide at position 16, introduced to prevent nonenzymatic spontaneous cleavage between C16 and A17^96,97^. For each reaction, 10 pmol of RNA was folded in buffer containing 300 mM NaCl, 10 mM Tris-HCl, 5 mM MgCl_2_ pH 8.0. Briefly the appropriate amount of RNA was diluted in H_2_O and heated 3 min in 80 °C followed by slow cooling. Then, at 50 °C a concentrated buffer was added to get final buffer solution and sample was continuously slowly cooled.

After 10 min incubation at 4 °C chemical mapping was conducted using two concentration of each reagent. To a 9 μl sample, 1 μl of 300 mM or 160 mM DMS in ethanol was added to give a final concentration of 30 or 15 mM DMS. For modification with CMCT, 9 μl of CMCT solution was added to the 9 μl of RNA sample. CMCT was diluted in a folding buffer to give a final concentration of 250, and 100 mM in the reaction mixture. Kethoxal was diluted in ethanol/water (1:3 v/v) to give a final concentration of 160 and 80 mM. After modification with kethoxal, 3 µl of 35 mM potassium borate solution was added to stabilize products of modification. Chemical modification reactions were incubated for 1.5 h at 4 °C. Reactions were stopped by precipitation with ethanol. The chemical modification reactions were repeated for a total of two replicates of each agent. The RNA in control reactions was treated the same, except no chemical reagents was added.

Modification sites were identified by primer extension. The DNA primer for reverse transcription (RT) was synthesized with 6-fluorescein (FAM) on the 5′ end (5’FAMGTTCCGGACCGAAGCCCG). The DNA primer was complementary to 3’ end of RNA (the cassette part). For each reverse transcription reaction, 10 pmol of primer was used. Primer extension was performed at 55 °C with SuperScript III reverse transcriptase using Invitrogen’s protocol. Reactions were stopped by addition of loading buffer containing urea and 10 mM EDTA, then chilling on ice. Prior to separation and read-out of cDNA products the samples were heated for 5 min at 95 °C and then separated on a 12% polyacrylamide denaturing gel (Figure S5).

The gel image from the Phosphorimager was analyzed using SAFA program to quantify nucleotide reactivities^98^. cDNA products were identified by comparing to sequencing lanes and to control lanes and the raw results from SAFA were normalized. To quantify chemical modification at each nucleotide, we first corrected for the background by subtracting the volume integral of the band in the control lane from the volume integral of experimental lane. For each of two experiments for each modification agent and each sequence, we characterized the modification extent by quartiles. When a nucleotide was in the highest quartile of RT stops in both experiments, we report the mapping as strong (Figure 4A). When a nucleotide was in the second highest quartile in both experiments or the highest quartile in one and the second highest quartile in the other, we report the cleavage as moderate.

### Transcriptome-Wide Calculations

We downloaded the set of m^6^A positions reported in the human transcriptome by Schwartz et al.^55^, which was available as their Supplementary Table S2. Using a Python program, for each entry for the human genome of “high confidence category” and with a RefSeq entry, we fetched the sequence from RefSeq^99^ using the Bio.Entrez module from Biopython^100^. To identify the exact position of the m^6^A in the transcript, we used the provided hg19 coordinates to identify the A in one of the expected sequence motifs (GGACA, GGACT, GGACC, GAACT, AGACT, AGACA, or TGACT) using the twobitreader Python package^101^ and the hg19 sequence downloaded from the UCSC genome browser^102^. Once the motif was identified in the genome, the sequence was found in the RefSeq sequence, and an 800 nucleotide FASTA sequence was generate with the m^6^A position at the 401^st^ position. For sequences in which the m^6^A was too close to the 5’ end or 3’ end to be in the 401^st^ position, up to 800 nucleotides were extracted with the m^6^A position at either the 5’ end or 3’ end. Sequences were generated with both A and 6 at the m^6^A position. In total, 18,155 high confidence m^6^A sites were found.

Next, the partition function was calculated for each 800 nucleotide sequence using the partition program from RNAstructure^44^. To determine the probability that the m^6^A position was buried in a helix, a custom C++ program was written using inheritance of the RNA class^44^. The probability of the i^th^ nucleotide being buried in a helix is the sum for all j of the probability the i-j base pair is sandwiched between the base pairs (i-1)-(j+1) and (i+1)-(j-1). Each of these can be determined using the partition function, Q, as a normalization factor and partial partition function for interior and exterior fragments:

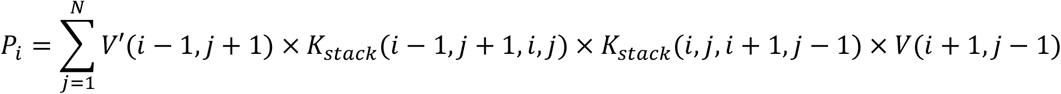

where P_i_ is the probability that nucleotide i is buried in a helix, N is the length of the sequence, V’(i,j) is the partition function for the exterior fragment of nucleotides 1 to i to j and to N given that i is paired to j, K_stack_(i,j,i+1,j-1) is the equilibrium constant for the base pair stack of base pairs i-j and (i+1)-(j-1), and V(i,j) is the partition function for the interior fragment from nucleotides i to j given that i is paired to j. Figure S6 diagrams V’ and V. These arrays of partition functions for sequence fragments are also explained in a description of the partition function calculation^89^.

### PARS calculations

To calculate PARS scores for human transcripts, we downloaded the dataset deposited by Wan et al.^56^ to the NCBI GEO (Gene Expression Omnibus)^103^. We used the mapped reads available for S1-treated (accessions GSM1226157, GSM1226159, and GSM1226161) and V1-treated (accessions GSM1226158, GSM1226160, and GSM1226162) samples. We calculated the PARS-score using^56^:

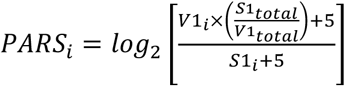

where PARS_i_ is the PARS score for the i^th^ nucleotide, V1_i_ is the number of reads in the V1-treated samples attributed to the i^th^ nucleotide, S1_i_ is the number of reads in the S1-treated samples attributed to the i^th^ nucleotide, S1_total_ is the total number of S1-treated sample reads, and V1_total_ is the total number of V1-treated sample reads. The ratio of S1_total_ and V1_total_ is a normalization factor. The addition of 5 in the numerator and denominator is a pseudocount to reduce the magnitude of scores for positions with few reads^56^. In total, entries were found for 18,026 transcripts of the 18,155 high-confidence m^6^A-containg transcripts found.

## Author Contributions

E.K. designed experiments, synthesized strands, performed optical melting experiments, performed chemical mapping experiments, and revised the manuscript, X.Z. contributed code to RNAstructure, R.M.W. contributed code to RNAstructure, R.K. synthesized phosphoramidites and strands and revised the manuscript, D.H.M. designed experiments, contributed code to RNAstructure, fit the nearest neighbor parameters, and drafted the manuscript.

