## Supplement for "Secondary Structure Prediction for RNA Sequences Including N^6^-methyladenosine"

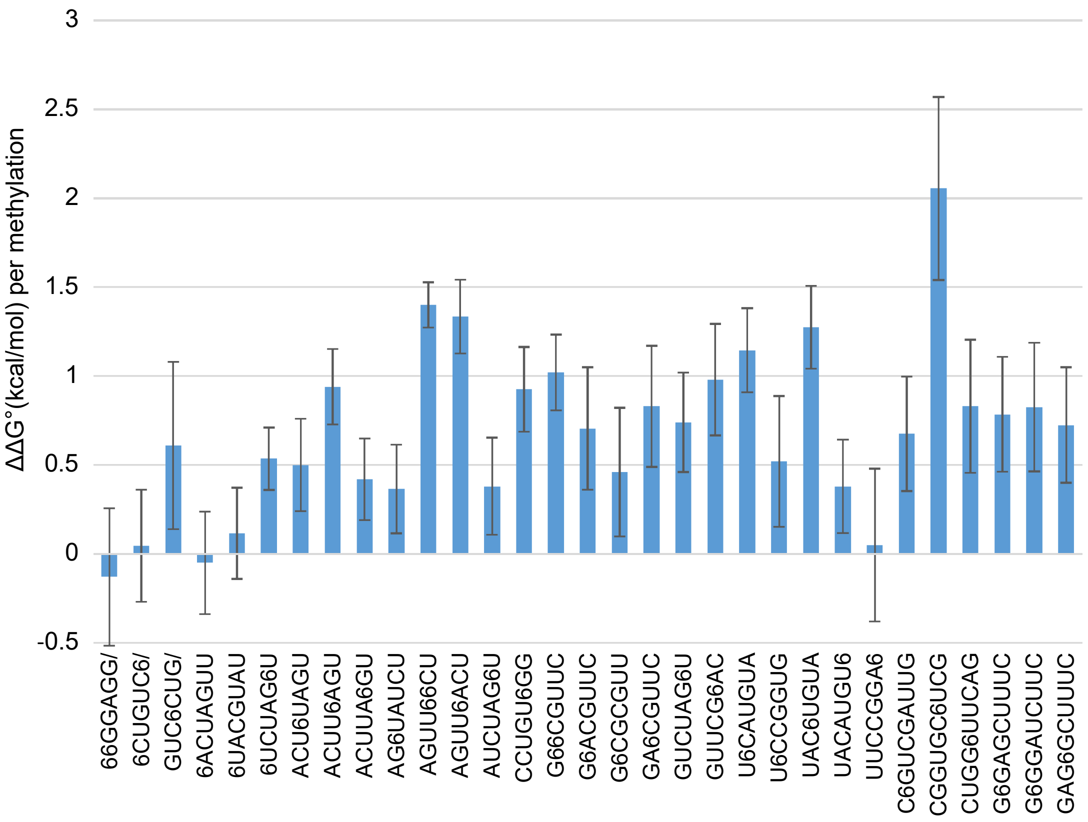


**Figure S1. The $\Delta\Delta G°_{37}$ for methylation is sequence dependent.** This plot shows the difference in stability between methylated and unmethylated duplexes, where the positive $\Delta\Delta G°_{37}$ means that the methylated duplex is less stable. 6 indicates the position of $m^6A$, and sequences ending with / are paired with a complementary strand. The stabilities of the unmethylated duplexes were derived from prior reports[37,51,73,74,104,105]. Uncertainty is estimated as 4% of the magnitude of free energy from each experiment, as estimated by Xia et al.[51]. The uncertainty in $\Delta\Delta G°_{37}$ derives from error propagation[82].

```
#Description of RNA Folding Alphabet including N6-methyadenosine
(m6A).

#The Bases field defines the characters allowed in the alphabet.
Bases
X = N = x = n
A = a
C = c
G = g
U = u = T = t
M = m = 6
I

#The Pairing field defines bases that can form canonical pairs.
Pairing
A U
G C
G U
M U

#The Single field defines bases that are not allowed to pair.
Single
a
c
g
u
t
m

#The non-interacting field indicates nucleotides that neither pair nor
stack.
Non-interacting
X

#The linker field indicates a special "nucleotide" that indicates a
break in the backbone.
Linker
I
```

**Figure S2. The RNAstructure Configuration File for the m$^6$A Alphabet.** The configuration indicates the nucleotides that are allowed and is read during the initialization of each program. Comments start with "#". The "Bases" field lists the available nucleotides. N or X can be used for bases that are not allowed to pair and provide no stacking stability. U or T can be used for uracil. M or 6 are used to indicate m$^6$A. I is an intermolecular linker that is used between two stands in bimolecular folding[106]. The "Pairing" field lists the canonical base pairs. "Single" lists nucleotides that cannot form pairs. In RNAstructure, nucleotides in lowercase are not allowed to base pair. "Non-interacting" specifies that N and X provide no stacking stability. Because they are equivalent according to "Bases", only X needs to be specified. "Linker" specifies that "I" is the intermolecular linker, and not an actual part of the sequence.

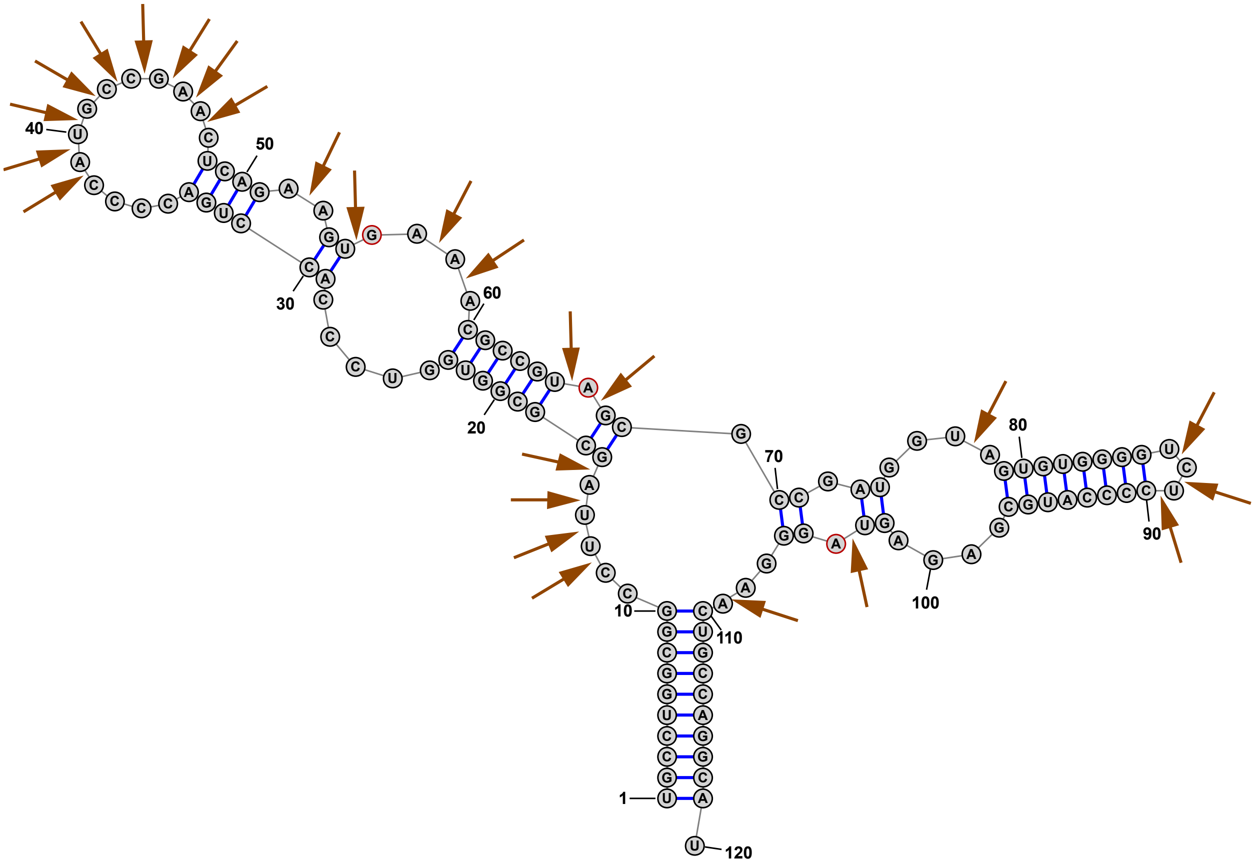


**Figure S3. S1 Nuclease Mapping of 5S rRNA.** This shows the data of Speek and Lind[61] superimposed upon the accepted structure of the *E. coli* 5S rRNA[107]. Arrows indicate backbone cleavages by S1 nuclease. The mapping pattern is consistent with cleavages both 5' and 3' to unpaired nucleotides. Nucleotides G56, A66, and A104 are unpaired bases with 5' cleavages where the 5' base is paired. This demonstrates that a subset of PARS S1 cleavages attributed to the 5' base being unpaired should be attributed to the 3' base being unpaired.

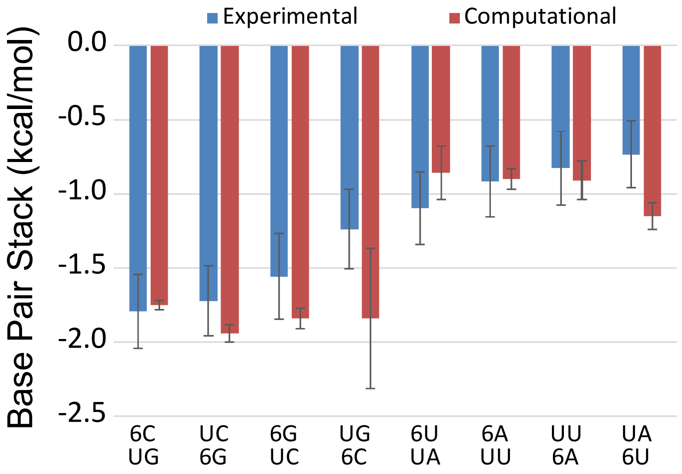


Figure S4. Comparison of Computationally-Estimated Stacking Nearest Neighbor Parameters (red)[66] to Our Experimental Nearest Neighbors (Blue). Overall the agreement of the estimates is excellent. The U-m$^6$A pair followed by a G-C pair has the largest deviation of 0.60 kcal/mol.

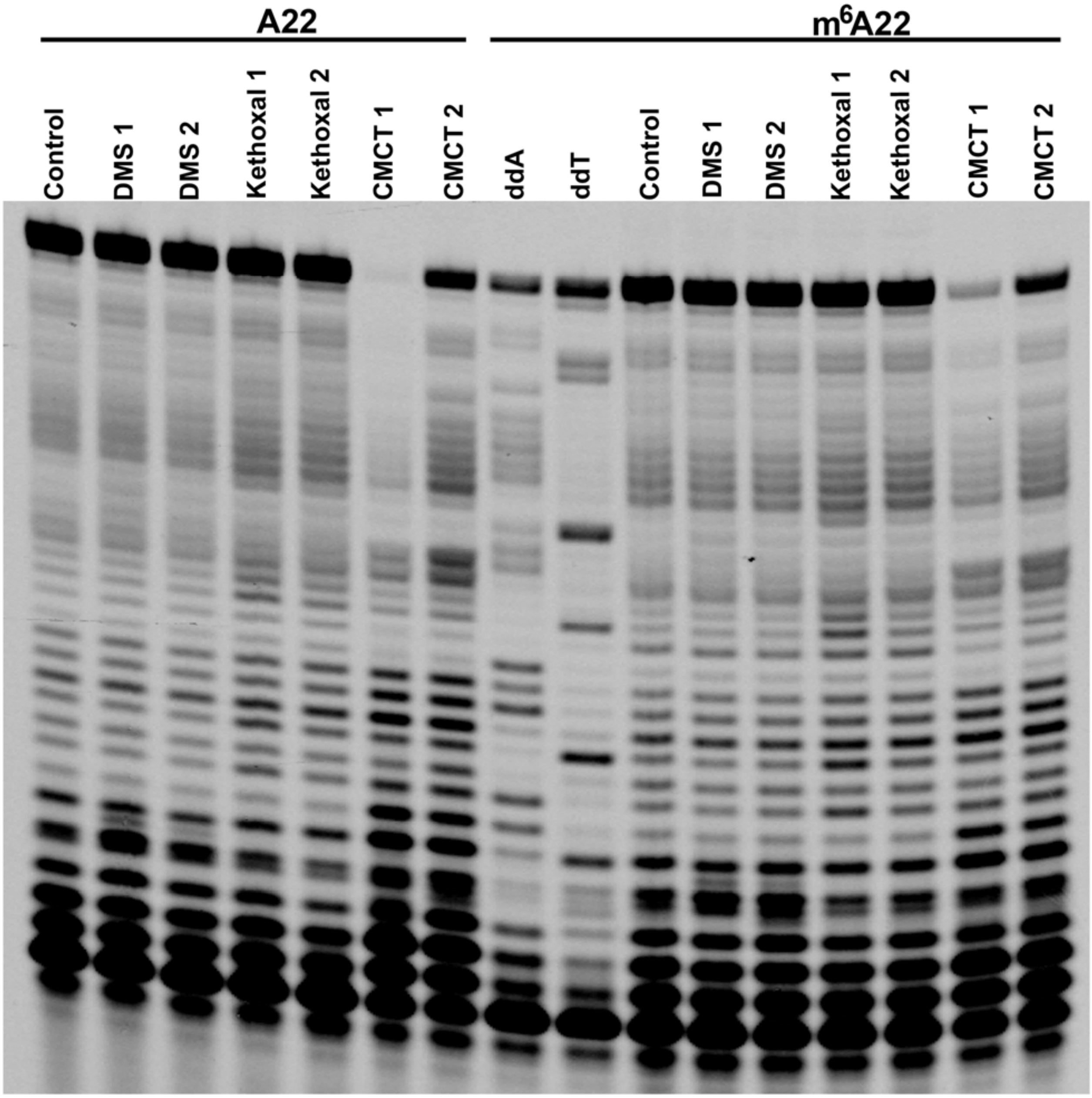


**Figure S5. Chemical Mapping Gel Image.**

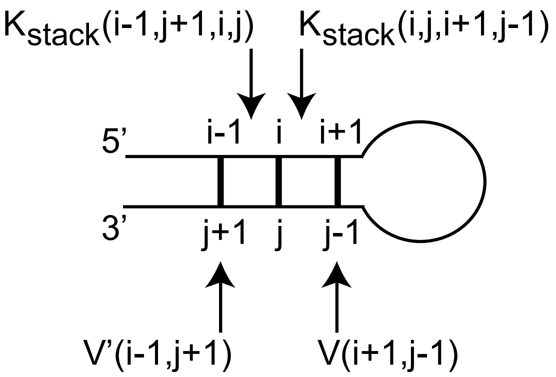


**Figure S6. The Calculation that the i-j Base Pair is Buried in a Helix.** In the illustration, the base pair i-j is stacking between adjacent pairs, (i+1)-(j-1) and (i-1)-(j+1). The contribution to the partition function for this configuration is the product of four terms: $K_{stack}$(i-1,j+1,i,j), the equilibrium constant for the stack of two base pairs; $K_{stack}$(i,j,i+1,j-1), the equilibrium constant for the other base pair stack; V(i+1,j-1), the partition function from nucleotides i+1 to j-1, inclusive, given that i+1 and j-1 base pair; and V'(i-1,j+1), the partition function from nucleotides 1 to i and also j to N (the length of the sequence), given that i-1 and j+1 base pair. The calculations for V and V' are provided by the recursions in Mathews[89], and follow the algorithm of McCaskill[47]. The total probability that nucleotide i is in a base pair that is stacked on adjacent base pairs in determined by summing these probabilities for all j to which nucleotide i can base pair.

**Table S1. Helical Duplexes Used to Fit Stacking Nearest Neighbor Parameters.** This table provides the results of the two-state fits to optical melting data. If the enthalpy changes of the average of the curve fits and the analysis of $T_M$ dependence agree to within 15%, the melting is consistent with two-state behavior.

| | Analysis of $T_M$ Dependence | | | Average of the Curve Fits | | |
|---|---|---|---|---|---|---|
| Sequence†: | $\Delta G°_{37}$ (kcal/mol) | ΔH° (kcal/mol) | ΔS° (cal $mol^{-1}$ $K^{-1}$) | $\Delta G°_{37}$ (kcal/mol) | ΔH° (kcal/mol) | ΔS° (cal $mol^{-1}$ $K^{-1}$) |
| 66GGAGG/ UUCCUCC | -9.80±0.28 | -61.3±7.0 | -166.3±22.1 | -9.75±0.42 | -55.2±8.5 | -146.5±26.5 |
| 6CUGUC6/ UGACAGU | -7.83±0.01 | -46.6±1.4 | -125.0±4.5 | -7.87±0.08 | -51.1±2.1 | -139.5±6.6 |
| GUC6CUG/ CAGUGAC | -8.01±0.04 | -61.0±2.4 | -170.8±7.8 | 8.06±0.12 | -61.8±4.5 | 173.3±14.4 |
| $6ACUAGUU_2$ | -7.26±0.10 | -54.7±4.5 | -153.0±14.3 | -7.23±0.16 | -51.1±2.2 | -141.7±7.0 |
| $6UACGUAU_2$ | -6.30±0.04 | -40.9±1.6 | -111.8±5.5 | -6.31±0.20 | -45.3±4.9 | -125.7±16.1 |
| $6UCUAG6U_2$ | -5.06±7.7 | -52.3±2.3 | -152.3±7.7 | -5.08±0.05 | -52.5±3.0 | -153.0±10.0 |
| $ACU6UAGU_2$ | -5.98±0.20 | -38.0±4.5 | -103.2±14.9 | -5.96±0.20 | -43.6±5.4 | -121.5±16.9 |
| $ACUU6AGU_2$ | -4.28±0.21 | -45.1±4.5 | -131.7±15.2 | -4.32±3.7 | -45.3±3.7 | -132.1±12.3 |
| $ACUUA6GU_2$ | -5.32±0.08 | -49.1±2.7 | -141.3±8.9 | -5.36±0.24 | -50.8±4.6 | -146.6±15.6 |
| $AG6UAUCU_2$ | -5.85±0.08 | -47.4±2.2 | -134.1±7.4 | -5.71±0.10 | -52.5±1.8 | -151.1±5.6 |
| $AGUU66CU_2$ | -0.76±0.41 | -54.8±4.0 | 174.5±14.2 | -1.96±0.67 | -43.5±5.6 | -134.0±20.2 |
| $AGUU6ACU_2$ | -3.69±0.25 | -45.3±3.7 | -134.1±12.7 | -3.95±0.22 | -42.0±4.0 | -122.9±13.4 |
| $AUCUAG6U_2$ | -6.44±0.04 | -56.9±3.4 | -162.7±11.0 | -6.42±0.16 | -50.5±3.0 | -142.2±9.6 |
| $CCUGU6GG_2$ | -4.96±0.17 | -49.8±4.1 | -144.6±13.7 | -4.93±0.12 | -53.7±7.6 | -157.5±24.7 |
| $G66CGUUC_2$ | -5.22±0.15 | -54.9±4.8 | -160.3±15.8 | -5.28±0.10 | -51.8±6.1 | -150.1±19.8 |
| $G6ACGUUC_2$ | -7.89±0.08 | -75.9±3.9 | -219.4±12.3 | -7.68±0.16 | -63.8±5.1 | -181.2±16.7 |
| $G6CGCGUU_2$ | -8.58±0.04 | -61.1±1.1 | -169.5±3.6 | -8.40±0.16 | -56.8±3.6 | -156.1±11.3 |
| $GA6CGUUC_2$ | -7.64±0.06 | -57.4±2.9 | -160.5±8.1 | -7.85±0.14 | -66.4±3.8 | -188.7±12.1 |
| $GGUU6ACC_2$ | -6.64±0.06 | -51.7±3.4 | -145.3±11.1 | -6.63±0.08 | -55.6±3.8 | -158.1±12.2 |
| $GUCUAG6U_2$ | -6.22±0.02 | -52.6±1.7 | -149.6±5.8 | -6.24±0.08 | -54.5±2.6 | -155.6±8.4 |
| $GUUCG6AC_2$ | -6.80±0.01 | -61.0±1.8 | -174.9±5.7 | -6.83±0.09 | -62.0±2.5 | -178.1±8.1 |
| $U6CAUGUA_2$‡ | -4.67±0.04 | -55.7±1.3 | -164.6±4.2 | 4.75±0.13 | 54.8±3.8 | 161.4±12.5 |
| $U6CCGGUG_2$ | -8.66±0.06 | -49.9±1.6 | -133.0±5.1 | -8.68±0.08 | -49.5±3.8 | -131.5±12.4 |
| $UAC6UGUA_2$‡ | -4.41±0.07 | -57.2±1.6 | -170.2±5.4 | 4.36±0.24 | 58.9±5.5 | 175.8±18.3 |
| $UACAUGU6_2$‡ | -6.20±0.01 | -57.5±1.4 | -165.4±4.6 | 6.27±0.05 | 56.8±5.1 | 163.0±16.5 |
| $UUCCGGA6_2$ | -10.69±0.13 | -65.6±2.0 | -177.1±6.2 | -10.58±0.18 | -172.5±7.6 | -64.1±2.5 |
| $C6GUCGAUUG_2$ | -7.35±0.02 | -58.9±1.9 | -166.3±6.2 | -7.37±0.08 | -60.9±1.4 | -172.7±5.0 |
| $CGGUGC6UCG_2$ | -10.65±0.11 | -86.0±2.5 | -243.2±7.8 | -10.60±0.12 | -84.5±2.2 | -238.5±6.8 |
| $CUGG6UUCAG_2$ | -8.49±0.18 | -77.4±7.0 | -222.2±22.0 | -8.38±0.07 | -72.3±1.7 | -205.0±5.4 |
| $G6GAGCUUUC_2$ | -7.25±0.01 | -63.7±1.5 | -182.0±4.8 | -7.23±0.06 | -62.8±1.5 | -179.2±4.7 |
| $G6GGAUCUUC_2$ | -8.18±0.11 | -66.8±4.1 | -189.1±13.0 | -8.14±0.11 | -65.7±2.0 | -185.6±6.3 |
| $GAG6GCUUUC_2$ | -7.37±0.07 | -76.8±4.6 | -223.7±14.7 | -7.39±0.14 | -77.9±1.8 | -227.2±5.9 |

† $m^6A$ is indicated as 6.

‡Reported previously[37].

**Table S2. Nearest Neighbor Stacks for m$^6$A-U pairs.** M$^6$A is indicated with 6.

| Nearest Neighbor Stack: | ΔG°$_{37}$ (kcal/mol) |
|---|---|
| 5'6C3'<br>3'UG5' | -1.79±0.25 |
| 5'UC3'<br>3'6G5' | -1.72±0.24 |
| 5'6G3'<br>3'UC5' | -1.56±0.29 |
| 5'UG3'<br>3'6C5' | -1.24±0.27 |
| 5'6U3'<br>3'UA5' | -1.10±0.25 |
| 5'6A3'<br>3'UU5' | -0.92±0.24 |
| 5'UU3'<br>3'6A5' | -0.83±0.25 |
| 5'UA3'<br>3'6U5' | -0.73±0.23 |
| 5'6U3'<br>3'UG5' | -0.69±0.32 |
| 5'6U3'<br>3'U65' | -0.46±0.36 |
| 5'UG3'<br>3'6U5' | -0.32±0.33 |
| 5'UU3'<br>3'6G5' | -0.32±0.27 |
| 5'663'<br>3'UU5' | -0.21±0.34 |
| 5'6G3'<br>3'UU5' | -0.03±0.29 |
| 5'U63'<br>3'6U5' | +1.45±0.57 |

**Table S3. Stabilities of dangling end, terminal mismatch, and loop sequences with m$^6$A (shown as "6").** Unpaired nucleotides are underlined.

| | | Analysis of $T_M$ Dependence | | | Average of Curve Fits | | |
|---|---|---|---|---|---|---|---|
| Sequence: | Motif: | ΔG°$_{37}$ (kcal/mol) | ΔH° (kcal/mol) | ΔS° (cal mol$^{-1}$ K$^{-1}$) | ΔG°$_{37}$ (kcal/mol) | ΔH° (kcal/mol) | ΔS° (cal mol$^{-1}$ K$^{-1}$) |
| ACAUGU6 †,‡<br>6UGUACA | 3' Dangling End | -5.79±0.02 | -62.0±1.8 | -181.3±5.9 | | | |
| GCGC6 †<br>6CGCG | 3' Dangling End | -8.89±0.04 | -52.4±0.9 | -140.4±2.8 | -8.78±0.04 | -50.1±0.09 | -133.2±2.9 |
| CCGG6 †<br>6GGCC | 3' Dangling End | -7.77±0.07 | -51.9±1.8 | -142.2±5.7 | -7.73±0.22 | -49.5±4.4 | -134.8±13.6 |
| 6AUGAAU †<br>UACGUA6 | 5' Dangling End | -6.49±0.02 | -46.8±1.8 | -130.1±6.0 | -6.51±0.12 | -47.4±3.2 | -131.9±10.3 |
| 6GCGC6 †<br>6CGCG6 | Terminal Mismatch | -8.21±0.21 | -42.6±3.8 | -111.0±11.7 | -8.39±0.26 | -45.4±3.4 | -119.3±10.3 |
| 6UGCGCA6 †<br>6ACGCGU6 | Terminal Mismatch | -9.92±0.21 | -54.5±3.0 | -143.7±8.9 | -9.91±0.34 | -54.7±3.8 | -144.4±11.2 |
| 6UGCGC66 †<br>66CGCGU6 | Terminal Mismatch | -9.70±0.26 | -51.7±3.6 | -135.3±10.6 | -9.81±0.27 | -53.4±3.1 | -140.5±9.1 |
| 6UGCGCAG †<br>GACGCGU6 | Terminal Mismatch | -9.74±0.15 | -51.4±2.1 | -134.3±6.5 | -10.23±0.20 | -58.5±2.6 | -155.6±7.9 |
| CGGCC6 †<br>6CCGGC | Terminal Mismatch | -9.06±0.19 | -50.4±2.9 | -133.2±8.7 | -9.20±0.22 | -52.2±1.8 | -138.7±5.1 |
| 6GGCGCUG †<br>GUCGCGG6 | Terminal Mismatch | -10.86±0.49 | -68.6±6.4 | -186.2±19.0 | -10.24±0.27 | -60.4±3.2 | -161.8±9.7 |
| GAUGCAU6 †<br>6UACGUAG | Terminal Mismatch | -7.29±0.09 | -54.3±3.4 | -151.6±10.7 | -7.37±0.12 | -59.2±3.1 | -167.0±9.9 |
| GGCGU6AUAGCC | Hairpin Loop | N.A. [A] | N.A. [A] | N.A. [A] | -3.19±0.11 | -33.3±1.1 | -97.3±3.5 |
| GCGG6UGC †<br>CGU6GGCG | Internal Loop | -4.30±0.01 | -44.6±0.12 | -129.9±0.4 | -4.25±0.04 | -45.4±0.7 | -132.6±2.5 |

†These are self-complementary duplexes. Both strands in duplexes are shown to demonstrate the motifs.

‡ This 3' dangling end was previously measured[37].

A. Hairpin loops are unimolecular and therefore the $T_M$ is not concentration-dependent.

**Table S4. Loop motif stabilities where 6 indicates $m^6A$.** The $\Delta\Delta G°_{37}$ is the stability of the motif with $m^6A$ minus the stability of the analogous motif with A; therefore, $\Delta\Delta G°_{37} < 0$ indicates $m^6A$ is more stabilizing than an analogous A. The determination of the motif stability is detailed in the Methods.

| Motif Sequence: | Motif Type: | Reference $\Delta G°_{37}$ (kcal/mol) | Motif $m^6A$ $\Delta G°_{37}$ (kcal/mol) | Motif A $\Delta G°_{37}$ (kcal/mol) | Motif $\Delta\Delta G°_{37}$ (kcal/mol) |
|---|---|---|---|---|---|
| U6<br>A | 3' Dangling End | -5.28±0.30 [A] | -0.43±0.20 | -0.21±0.19 [B] | -0.22±0.27 |
| C6<br>G | 3' Dangling End | -4.61±0.18 [C] | -2.14±0.20 | -1.66±0.18 [D] | -0.49±0.27 |
| G6<br>C | 3' Dangling End | -4.55±-0.18 [E] | -1.61±0.18 | -1.15±0.16 [F] | -0.47±0.24 |
| 6A<br>U | 5' Dangling End | -4.42±0.33 [A] | -1.04±0.21 | -0.48±0.20 [G] | -0.56±0.29 |
| C6<br>G6 | Terminal Mismatch | -4.61±0.18 [C] | -1.80±0.18 | -1.52±0.18 [H] | -0.29±0.26 |
| A6<br>U6 | Terminal Mismatch | -8.22±0.33 [I] | -0.85±0.26 | -0.73±0.25 [J] | -0.12±0.36 |
| 66<br>U6 | Terminal Mismatch | -7.16±0.61 [A] | -1.27±0.36 | -0.73±0.36 [J] | -0.54±0.51 |
| AG<br>U6 | Terminal Mismatch | -8.22±0.33 [I] | -0.76±0.25 | -0.78±0.26 [J] | 0.02±0.36 |
| C6<br>GC | Terminal Mismatch | -5.37±0.21 [K] | -1.85±0.21 | -1.47±0.20 [H] | -0.38±0.29 |
| UG<br>G6 | Terminal Mismatch | -8.42±0.34 [L] | -1.22±0.27 | -0.48±0.25 [M] | -0.74±0.37 |
| U6<br>AG | Terminal Mismatch | -4.42±0.33 [A] | -1.44±0.22 | -1.23±0.21 | -0.18±0.31 |
| CGU6AUAG | Hairpin Loop | -6.68±0.11 [A] | 3.49±0.17 | 3.26±0.17 [N] | 0.23±0.24 |
| GG6U<br>U6GG | Internal Loop | -4.82±0.33 [O] | 0.52±0.37 | 0.19±0.38 [P] | 0.33±0.53 |

A. Estimated with nearest neighbor parameters[51].
B. Analogous A-containing duplex measured previously[37].
C. Reference was measured previously[108].
D. Analogous A-containing duplex measured previously[109].
E. Reference was measured previously[110].
F. Analogous A-containing duplex measured previously[110].
G. Analogous A-containing duplex measured previously[111].
H. Analogous A-containing duplex measured previously[112].
I. Reference was measured previously[109].
J. Analogous A-containing duplex measured previously[113].
K. Reference was measured previously[79].
L. Reference was measured previously[114].
M. Analogous A-containing duplex measured previously[115].
N. Analogous A-containing hairpin measured previously[63].
O. Reference duplex estimated with nearest neighbor parameters[73].
P. Analogous A-containing duplex measured previously[116].

**Table S5. Additional optical melting experiments to test nearest neighbor model.** 6 is used to represent $m^6A$.

| | Analysis of $T_M$ Dependence | | | Average of the Curve Fits | | |
|---|---|---|---|---|---|---|
| Sequence: | $\Delta G°_{37}$ (kcal/mol) | $\Delta H°$ (kcal/mol) | $\Delta S°$ (cal $mol^{-1}$ $K^{-1}$) | $\Delta G°_{37}$ (kcal/mol) | $\Delta H°$ (kcal/mol) | $\Delta S°$ (cal $mol^{-1}$ $K^{-1}$) |
| $GGACUAGUCC_2$ | -16.19±0.56 | -95.0±5.7 | -254.2±16.8 | -16.99±0.12 | -103.2±0.9 | -278.2±2.8 |
| $GG6CUAGUCC_2$ | -15.36±0.75 | -101.7±8.1 | -273.4±23.9 | -15.54±0.22 | -103.6±1.9 | -283.9±5.5 |
| $CGCGCG_2$ | -9.38±0.14 (9.11±0.05)† | -47.8±1.9 (54.4±0.5)† | -124.0±5.9 (146.2±1.6)† | -10.26±0.36 (9.06±0.14)† | -59.8±2.2 (53.1±2.0)† | -159.9±6.3 (-142.1±6.1)† |
| $CGCGCG6_2$ | -12.25±0.22 | -68.3±2.4 | -180.7±7.0 | -12.19±0.12 | -67.6±0.9 | -178.7±2.6 |

†Previous measurement of $CGCGCG_2$ with a 3’ terminal phosphate, measured by Freier et al.[108] and reported by SantaLucia et al.[117].

**Table S6. Accuracy of Nearest Neighbor Estimates.** 6 stands for $m^6A$.

| Sequence: | Experimental $\Delta G°_{37}$ (kcal/mol) | Estimated $\Delta G°_{37}$ (kcal/mol) | Error in Estimate ($\Delta\Delta G°_{37}$; kcal/mol) |
|---|---|---|---|
| $GGACUAGUCC_2$ | -16.19±0.56 | -16.67±0.35 | 0.48±0.73 |
| $GG6CUAGUCC_2$ | -15.36±0.75 | -14.51±0.76 | -0.85±0.97 |
| $CGCGCG_2$ | -9.38±0.14 | -9.40±0.38 | 0.02±0.54 |
| $CGCGCG6_2$ | -12.25±0.22 | -12.26±0.68† | 0.01±0.84 |

†The uncertainty estimate for the dangling A increment was taken from Zuber et al.[48].

**Table S7. The Stacking Nearest Neighbor Stability to be Fit and the Estimate of the Fit.** "$\Delta G°_{37}$ Stacks with m[6]A-U pairs" is the stability of the duplex (Table S1) minus the stability of Watson-Crick and G-U pair stacks and minus the stability of A-U end penalties, symmetry penalties, and initiation penalties. This is the term fit in the regression. The "$\Delta G°_{37}$ Estimate" is estimated from the fit nearest neighbor parameters (Table S2). "Residual" is the difference between the two. The regression minimizes the sum of squares of residuals.

| Sequence†: | $\Delta G°_{37}$ Stacks with $m^6$A-U pairs (kcal/mol) | $\Delta G°_{37}$ Estimate (kcal/mol) | Residual (kcal/mol) |
|---|---|---|---|
| 66GGAGG/ UUCCUCC | -2.94 | -1.76 | -1.18 |
| 6CUGUC6/ UGACAGU | -3.14 | -3.03 | -0.11 |
| GUC6CUG/ CAGUGAC | -3.32 | -3.03 | -0.29 |
| $6ACUAGUU_2$ | -1.81 | -1.83 | 0.02 |
| $6UACGUAU_2$ | -1.32 | -2.19 | 0.87 |
| $6UCUAG6U_2$ | -4.09 | -4.35 | 0.26 |
| $ACU6UAGU_2$ | -2.76 | -1.93 | -0.83 |
| $ACUU6AGU_2$ | -1.06 | -0.39 | -0.67 |
| $ACUUA6GU_2$ | -4.93 | -4.77 | -0.16 |
| $AG6UAUCU_2$ | -5.78 | -5.64 | -0.14 |
| $AGUU66CU_2$ | -2.02 | -2.55 | 0.53 |
| $AGUU6ACU_2$ | -0.47 | -0.39 | -0.08 |
| $AUCUAG6U_2$ | -6.36 | -5.64 | -0.73 |
| $CCUGU6GG_2$ | -3.68 | -3.75 | 0.07 |
| $G66CGUUC_2$ | -7.38 | -7.44 | 0.06 |
| $G6ACGUUC_2$ | -5.57 | -5.28 | -0.29 |
| $G6CGCGUU_2$ | -4.96 | -4.22 | -0.74 |
| $GA6CGUUC_2$ | -5.10 | -5.24 | 0.14 |
| $GGUU6ACC_2$ | -0.16 | -0.39 | 0.23 |
| $GUCUAG6U_2$ | -5.25 | -4.82 | -0.43 |
| $GUUCG6AC_2$ | -4.48 | -5.28 | 0.80 |
| $U6CAUGUA_2$ | -4.77 | -5.05 | 0.28 |
| $U6CCGGUG_2$ | -4.30 | -4.23 | -0.07 |
| $UAC6UGUA_2$ | -2.70 | -2.93 | 0.24 |
| $UACAUGU6_2$ | -0.92 | -1.47 | 0.55 |
| $UUCCGGA6_2$ | -1.63 | -1.65 | 0.02 |
| $C6GUCGAUUG_2$ | -3.01 | -2.54 | -0.47 |
| $CGGUGC6UCG_2$ | -3.43 | -3.86 | 0.43 |
| $CUGG6UUCAG_2$ | -1.03 | -1.09 | 0.06 |
| $G6GAGCUUUC_2$ | -3.17 | -3.50 | 0.33 |
| $G6GGAUCUUC_2$ | -3.36 | -3.50 | 0.14 |
| $GAG6GCUUUC_2$ | -3.07 | -3.75 | 0.68 |

Table S8 the number of occurrences of each stacking parameter in the set of fit helices.

| Nearest Neighbor Stack: | Occurrences in Fitting Set: |
|---|---|
| 5'6C3'<br>3'UG5' | 14 |
| 5'UC3'<br>3'6G5' | 18 |
| 5'6G3'<br>3'UC5' | 7 |
| 5'UG3'<br>3'6C5' | 8 |
| 5'6U3'<br>3'UA5' | 6 |
| 5'6A3'<br>3'UU5' | 12 |
| 5'UU3'<br>3'6A5' | 6 |
| 5'UA3'<br>3'6U5' | 6 |
| 5'6U3'<br>3'UG5' | 4 |
| 5'6U3'<br>3'U65' | 5 |
| 5'UG3'<br>3'6U5' | 4 |
| 5'UU3'<br>3'6G5' | 6 |
| 5'663'<br>3'UU5' | 5 |
| 5'6G3'<br>3'UU5' | 6 |
| 5'U63'<br>3'6U5' | 4 |
